# Constrained Evolutionary Design of Matrixyl Analogs: Balancing Permeability and Functional Preservation Through Computational Optimization

**DOI:** 10.64898/2026.05.12.724473

**Authors:** Nikolaos Komianos, Pranadarth Prakash

## Abstract

Matrixyl (palmitoyl pentapeptide-4, KTTKS core) is a collagen-stimulating peptide used in topical anti-ageing products, but its in-use efficacy is limited by poor permeation through the stratum corneum. We describe a deterministic computational workflow that combines a tournament genetic algorithm and NSGA-II with exact RDKit molecular descriptors to search the fixed-length, edit-distance-2 neighbourhood of KTTKS (3,706 candidate sequences) for analogs with descriptors more favourable for passive transdermal diffusion. The search returns a 9-member Pareto frontier that quantifies the trade-off between predicted permeability and motif preservation. Five of the nine frontier members carry the same substitution, lysine to proline at position 4 (K4P). This single change lowers the topological polar surface area by 25.6%, removes the +1 charge contributed by lysine, and reduces the functional-preservation score from 1.00 (KTTKS) to 0.67. The frontier ranking is unchanged by *±*30% perturbations to the TPSA and *M*_*w*_ penalty weights and by a 30% increase in the LogP penalty; only a 30% reduction in the LogP penalty produces rank movement. The frontier matches the ground-truth Pareto set obtained by exhaustive enumeration of all 3,706 candidates (precision and recall both 100%). On the basis of these results we recommend three sequences for experimental validation: PTTPS (largest predicted gain), KTTPS (single-mutation, conservative), and KTTPP (backup). All code, results, and figures are released under MIT and CC BY 4.0.

## 1 Introduction

### 1.1 Motivation and Problem Statement

Peptides are a growing class of pharmaceutical and cosmetic actives. The peptide therapeutics market alone was valued at roughly $40 billion in 2018 and has continued to grow [1]. Within cosmetic peptides, Matrixyl (palmitoyl pentapeptide-4, KTTKS core) is one of the longest-marketed actives in topical anti-ageing products and has shown collagen-stimulating activity in dermal fibroblast culture as well as measurable wrinkle improvement in a controlled clinical trial [2]. The in-use efficacy of topical Matrixyl is nevertheless limited by poor permeation through the stratum corneum, which is the dominant constraint on biologically active peptides delivered through the skin [3].

The barriers to topical peptide delivery are not unique to Matrixyl. Bioactive peptides typically have molecular weights (*M*_*w*_) in the 500–3,000 Da range, topological polar surface areas (TPSA) of 100–250 Å^2^, and net charges of ±2 to ±4, all of which violate the small-molecule transdermal rules established by Potts and Guy [4]. The standard industrial response is N-terminal lipidation, as in Pal-KTTKS, which improves LogP but increases molecular weight, with only modest net benefit for skin delivery [5].

Most peptide optimization workflows treat permeability as a downstream problem to be solved through formulation, e.g. with penetration enhancers or liposomes. In this work we treat topical delivery as a first-class objective, alongside motif preservation, and ask whether constrained evolutionary search can discover Matrixyl analogs with descriptors more favourable for passive transdermal diffusion while still resembling the parent KTTKS sequence.

### 1.2 Computational Approaches to Peptide Design

Protein language models (PLMs) such as ESM-2 [6] and structure prediction models such as AlphaFold [7] have substantially advanced protein design from sequence. PLMs capture sequence naturalness and have been used to generate functional protein sequences across diverse families [8], and diffusion-based generative models have been applied to backbone-level de novo protein design [11]. These approaches are well suited to long protein sequences, but they have two limitations for the present problem. First, they are trained predominantly on full-length proteins and may not capture the functional constraints of short cosmetic peptides. Second, they do not directly optimize for physicochemical descriptors such as TPSA or LogP, which determine passive transdermal diffusion.

Multi-objective evolutionary algorithms (MOEAs) take a complementary approach. NSGA-II [9] is the de facto standard MOEA and is widely used in cheminformatics; deep-learning approaches like the cross-docked CNN models of Francoeur et al. [10] typically supply scoring functions, but the optimization over a combinatorial sequence space is still well-suited to evolutionary search. An evolutionary approach also has the practical advantages that every objective function is written down explicitly, every constraint is interpretable, and the Pareto frontier shows the trade-off between objectives directly without a learned scalar utility.

For the Matrixyl problem the small search space (3,706 candidates within edit distance 2 of KTTKS) makes evolutionary search particularly attractive. The whole neighbourhood can be enumerated, so we can validate the evolved Pareto frontier against the ground-truth Pareto frontier rather than against a benchmark. Each mutation on the frontier can also be interpreted directly in terms of changes to the underlying descriptors (charge, hydrogen-bond count, polar surface area), which makes the resulting candidates easier to triage for experimental validation.

### 1.3 Research Questions and Hypotheses

We address three questions:

1. Can a constrained evolutionary search find Matrixyl-family analogs with descriptors more favourable for passive transdermal diffusion than the parent KTTKS?
2. Which substitutions, if any, recur on the Pareto frontier, and can they be interpreted in terms of the underlying descriptors?
3. Is the trade-off between predicted permeability and motif preservation stable to perturbations in the penalty weights of the scoring function?

Our prior expectations, set before the search was run, were that NSGA-II would return a Pareto frontier of roughly 5–15 candidates with a clear trade-off between predicted permeability and motif preservation, that mutations at the two lysine positions would dominate the frontier, and that the frontier ranking would be insensitive to ±30% changes in the penalty weights.

## 2 Methods

### 2.1 Baseline Peptides and Search Space Definition

#### 2.1.1 Reference Sequence

The Matrixyl core peptide, `KTTKS` (Lys-Thr-Thr-Lys-Ser), was selected as the reference sequence. This sequence represents the bioactive cargo of the commercially successful palmitoyl pentapeptide-4 (PubChem CID 9897237). The unmodified pentapeptide has the following documented properties:

- Molecular weight: 563.65 Da (calculated via RDKit)
- Net charge at pH 7.4: +2 (two lysine residues, deprotonated C-terminus)
- Canonical amino acids: Yes

#### 2.1.2 Search Space Constraints

To maintain biological plausibility and tractability, the search space was constrained as follows:

- **Backbone length**: Fixed at 5 residues (no insertions or deletions in main experiments).
- **Amino acid alphabet**: Canonical L-amino acids only (A, C, D, E, F, G, H, I, K, L, M, N, P, Q, R, S, T, V, W, Y).
- **Edit distance**: Maximum Levenshtein distance of 2 (ED≤ 2) from the reference KTTKS. This constraint was chosen to balance explorability with motif recognizability; candidates at ED= 2 differ by exactly 2 single-residue substitutions.
- **Search space size**: The fixed-length ED≤ 2 neighborhood enumerates to exactly 3,706 unique valid sequences (verified via dynamic-programming enumeration).

#### 2.1.3 Control Molecules

For validation and benchmarking, the following control molecules were evaluated through the same scoring pipeline:

- **KTTKS**: Unmodified Matrixyl core (baseline reference).
- **Pal-KTTKS**: N-terminal palmitoyl-conjugated KTTKS (*M*_*w*_ 802.07 Da, SMILES from Pub-Chem CID 9897237).
- **GPKGDPGA**: Collagen-like control motif (previously used in early iterations; included for completeness).

### 2.2 Molecular Descriptor Calculations

All molecular properties were calculated using RDKit 2023.09.1, ensuring exact, reproducible results independent of heuristic approximations.

#### 2.2.1 Peptide Structure Construction

For each canonical amino-acid sequence, a molecular structure was constructed as follows:

1. Convert the sequence to a linear peptide backbone with standard N-terminus (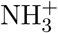 at pH 7.4) and C-terminus (COO^−^ at pH 7.4).
2. Attach side chains for each residue at the appropriate stereochemistry (all L-configuration).
3. For lipidated sequences (Pal-KTTKS), attach a 16-carbon palmitoyl chain via amide bond to the N-terminal amino group.
4. Sanitize and validate the resulting molecular structure using RDKit standard routines.

#### 2.2.2 Descriptor Suite

The seven molecular descriptors used to quantify topical-delivery character are summarized in Table 1.

**Table 1:**
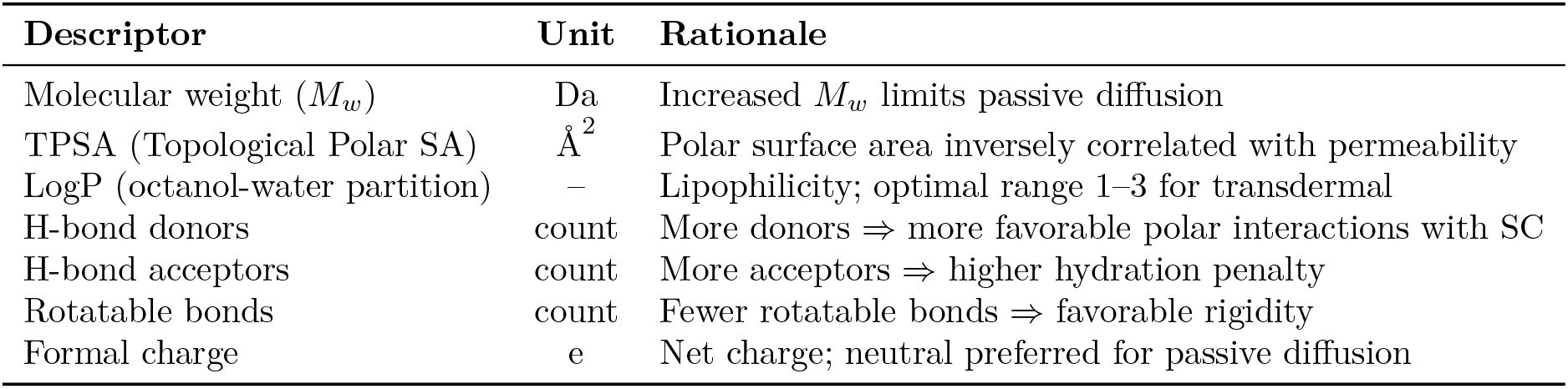
Molecular descriptors used in the penetration score, with units and the physico-chemical rationale for each.

| Descriptor | Unit | Rationale |
| --- | --- | --- |
| Molecular weight ($M_w$) | Da | Increased $M_w$ limits passive diffusion |
| TPSA (Topological Polar SA) | $\text{\AA}^2$ | Polar surface area inversely correlated with permeability |
| LogP (octanol-water partition) | – | Lipophilicity; optimal range 1–3 for transdermal |
| H-bond donors | count | More donors ⇒ more favorable polar interactions with SC |
| H-bond acceptors | count | More acceptors ⇒ higher hydration penalty |
| Rotatable bonds | count | Fewer rotatable bonds ⇒ favorable rigidity |
| Formal charge | e | Net charge; neutral preferred for passive diffusion |

**Table 2:**
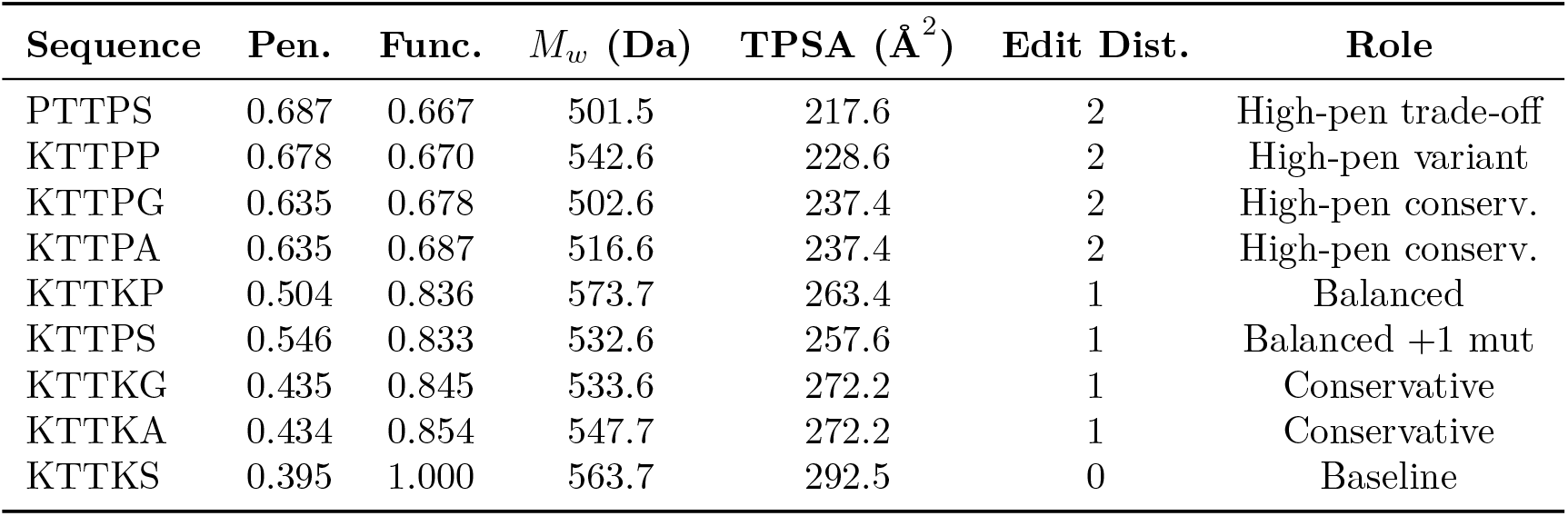
Pareto frontier: 9 candidates from NSGA-II multi-objective optimization.

**Table 3:**
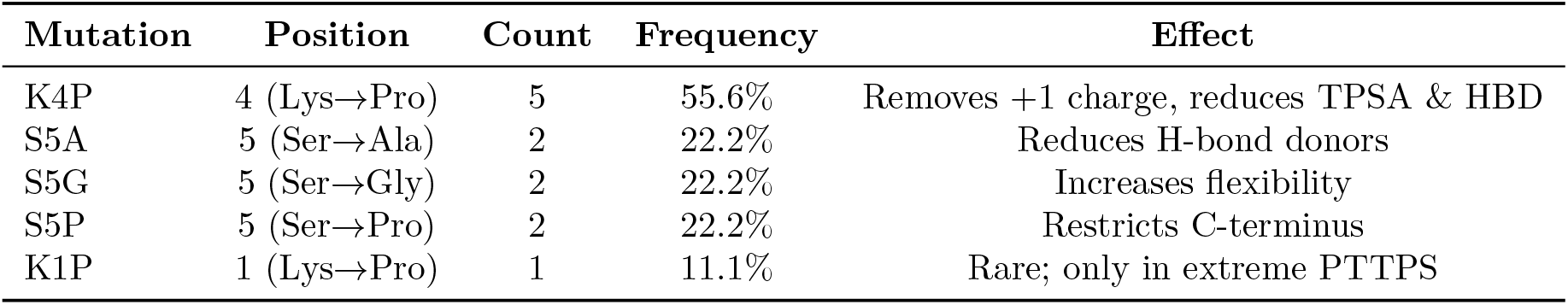
Mutation enrichment on the Pareto frontier (e.g., K4P denotes a Lys →Pro substitution at position 4). Counts and frequencies are computed across the 9-member frontier.

**Table 4:**
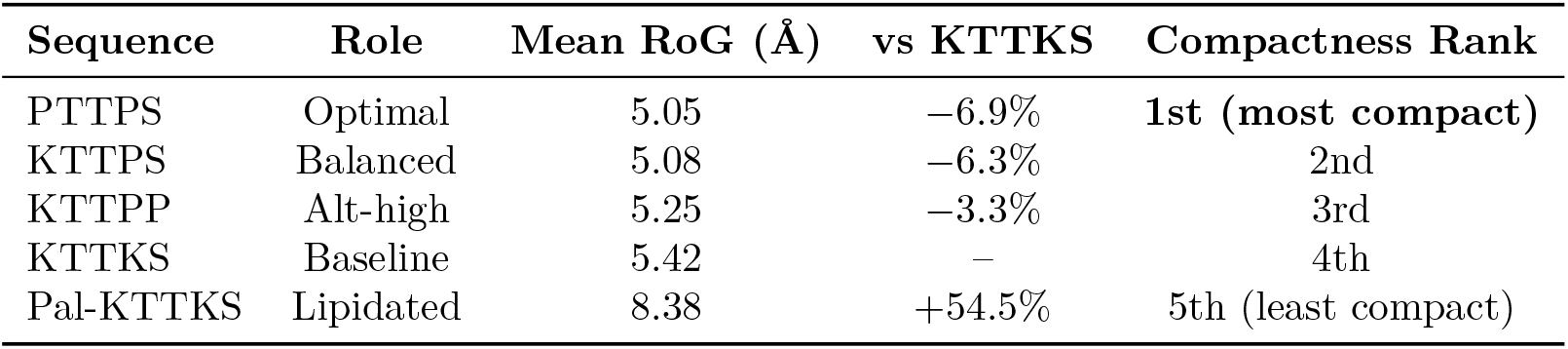
Radius of gyration (RoG), a one-number summary of molecular extent computed from RDKit ETKDG ensembles (Eq. 11): 8 conformers per unmodified peptide; 3 for Pal-KTTKS, whose flexible palmitoyl tail dominates the conformational extent. Lower RoG indicates a more compact, diffusion-favorable conformation.

**Table 5:** Extended search space at edit distance 3. The two “penetration gain vs baseline” values are the gain of each frontier’s best candidate against the unmodified KTTKS baseline (PTTPS for ED ≤ 2, KFFKP for ED ≤ 3). The ED ≤ 2 → ED ≤ 3 delta in the best candidate is +14.2% (0.687 → 0.785); see body text.

| Metric | $ED \leq 2$ | $ED \leq 3$ |
| --- | --- | --- |
| Frontier size | 9 | 20 |
| Best penetration | 0.687 (PTTPS) | 0.785 (KFFKP) |
| Penetration gain vs baseline | +74% | +99% |
| All $ED \leq 2$ frontier members on $ED \leq 3$ frontier | – | 9/9 (100%) |

**Table 6:** Ranking stability of the 9-member Pareto frontier under ±30% perturbations of the per-descriptor penalty weights. Rank change is the absolute change in a candidate’s position relative to the unperturbed baseline ranking. Five of the six perturbations leave the ranking completely unchanged; only a 30% reduction in the LogP penalty produces movement.

| Parameter Variation | Mean Rank Change | Max Rank Change |
| --- | --- | --- |
| TPSA $-30\%$ | 0.00 | 0 |
| TPSA $+30\%$ | 0.00 | 0 |
| $M_w$ $-30\%$ | 0.00 | 0 |
| $M_w$ $+30\%$ | 0.00 | 0 |
| LogP $-30\%$ | 1.78 | 6 |
| LogP $+30\%$ | 0.00 | 0 |
| Mean across 6 scenarios | 0.30 | 1.00 |

**Table 7:**
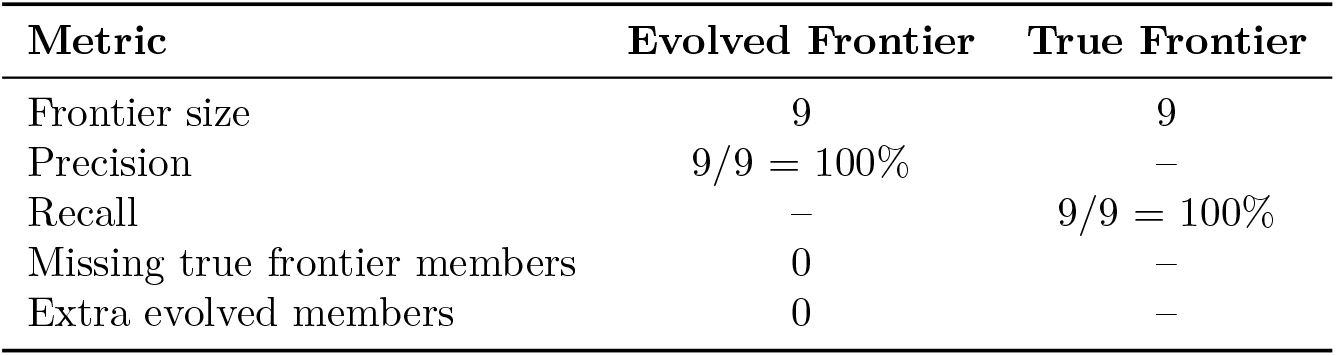
Exhaustive enumeration validation: NSGA-II recovered the true Pareto frontier with perfect precision and recall.

| Metric | Evolved Frontier | True Frontier |
| --- | --- | --- |
| Frontier size | 9 | 9 |
| Precision | 9/9 = 100% | – |
| Recall | – | 9/9 = 100% |
| Missing true frontier members | 0 | – |
| Extra evolved members | 0 | – |

All descriptors were calculated using standard RDKit definitions and cross-validated against PubChem data for known molecules (e.g., Pal-KTTKS against CID 9897237).

### 2.3 Objective Function Design

#### 2.3.1 Penetration Score

The penetration score is a composite normalized metric (range [0, 1]) that penalizes deviations from optimal delivery-favorable ranges:

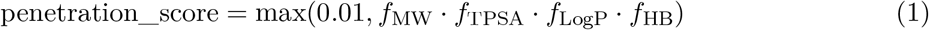

where:

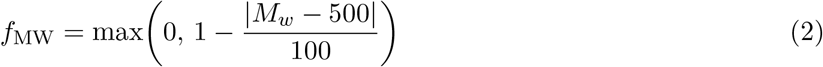

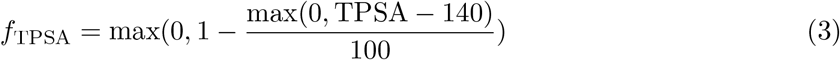

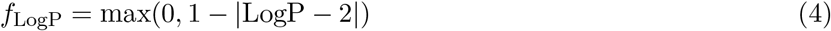

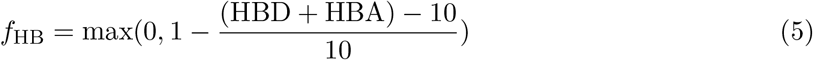

##### Rationale

The thresholds (*M*_*w*_ < 500 Da, TPSA < 140 Å^2^, LogP ≈ 1–3) derive from extensive transdermal drug permeability literature [4, 12]. We apply these as *soft penalties* (not hard cutoffs) because peptides often violate these rules and still permeate, especially when formulated. The multiplicative structure of Eq. 1 ensures that failure to meet any single objective significantly reduces overall score.

#### 2.3.2 Functional Preservation Score

The functional preservation score (range [0, 1]) quantifies how much a candidate resembles the KTTKS motif using a weighted combination of sequence identity, edit distance, and conservative substitution similarity:

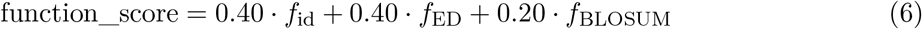

where:

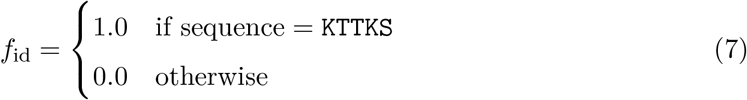

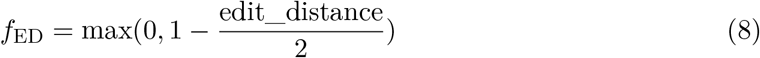

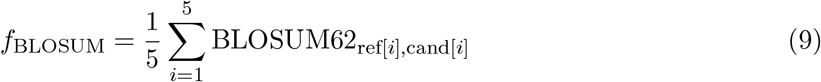

The BLOSUM62 score is normalized by dividing the sum by 5 (sequence length) and then by the maximum possible BLOSUM62 value (4.0 for identical residues), yielding a score in [0, 1].

##### Rationale

Position 4 lysine is known to be critical for collagen-stimulating activity in matrikines [13]. The weighted scheme of Eq. 6 reflects uncertainty: exact identity is valued (0.40 weight) but edit distance also matters (0.40 weight), and conservative substitutions should not be penalized as heavily as drastic mutations (0.20 weight).

#### 2.3.3 Synthesis Feasibility Score

For the initial fixed-length ED≤ 2 search space using only canonical amino acids, the synthesis feasibility score is constant:

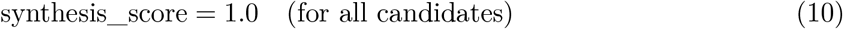

This score was defined to allow future extension to non-canonical residues or modified backbones, where synthesis complexity could be explicitly penalized.

### 2.4 Evolutionary Algorithms

#### 2.4.1 Phase 1: Tournament Selection (Single-Objective)

**Algorithm**: Generational genetic algorithm with tournament selection and elitism.

##### Parameters

- Population size: 100 individuals
- Generations: 100
- Tournament size: 3 (select 3 random individuals per tournament, pick best)
- Mutation probability: 0.2 per individual per generation
- Elite count: 2 (preserve best 2 individuals unchanged each generation)
- Random seed: 42 (deterministic reproducibility)

##### Operators

- **Mutation**: Select one position uniformly at random; replace with a uniformly random canonical amino acid.
- **Crossover**: Disabled (mutation-only evolution is simpler to interpret for short peptides).
- **Repair**: All offspring are validated against SearchSpaceRules (edit distance, alphabet, length). Invalid mutations are replaced by resampling.
- **Objective**: Maximize penetration_score × functional_preservation_score (scalar weighted objective).

#### 2.4.2 Phase 2: NSGA-II (Multi-Objective)

**Algorithm**: Non-dominated sorting genetic algorithm II (local reimplementation without external dependencies).

##### Parameters

- Population size: 100 individuals
- Generations: 100
- Tournament size: 2
- Mutation probability: 0.2 per individual
- Random seed: 42

##### Objectives (all maximized)

1. Penetration score
2. Functional preservation score
3. Synthesis feasibility score (constant: 1.0)

##### Selection

Binary tournament on (Pareto rank, crowding distance). Higher rank is worse (deeper into dominated set); same rank selects on crowding distance (prefer diverse solutions).

##### Survival

Elitist environmental selection. Merge parent population (100) and offspring (100 from mutation-only reproduction) into a pool of 200. Rank by non-dominance, then by crowding distance. Select top 100 for next generation. Deduplicate by sequence before final frontier computation.

##### Pareto Frontier Definition

A solution is non-dominated (on the frontier) if no other solution dominates it on all three objectives simultaneously.

### 2.5 Validation Strategy

#### 2.5.1 Exhaustive Enumeration

Because the fixed-length ED≤ 2 search space is enumerable (3,706 sequences), we performed complete enumeration as ground truth:

1. Generate all 3,706 valid KTTKS-family sequences via dynamic-programming Levenshtein distance.
2. Evaluate each sequence using identical CandidateEvaluator (same descriptor calculations, same scoring functions).
3. Compute the true Pareto frontier via all-pairs dominance comparison.

The evolutionary frontier was then compared against the exhaustive frontier to compute:

- **Precision**: Fraction of evolved frontier members that are on the true frontier.
- **Recall**: Fraction of true frontier members found by the algorithm.

#### 2.5.2 Sensitivity Analysis

To assess robustness, Phase 2 was re-run with ±30% variations in penalty weights:

- TPSA penalty multiplied by 0.7, 1.0, 1.3
- *M*_*w*_ penalty multiplied by 0.7, 1.0, 1.3
- LogP penalty multiplied by 0.7, 1.0, 1.3

For each scenario, candidate rankings were recomputed and compared to the baseline ranking.

Metrics reported: mean rank change, max rank change, fraction of top-candidate stability.

#### 2.5.3 Extended Search Space

Edit distance 3 (ED≤ 3) was evaluated to test for diminishing returns:

- Search space size: ≈ 100k candidates (not exhaustively enumerated due to size)
- Algorithm: Phase 2 (NSGA-II) with identical parameters
- Comparison: Best penetration score, frontier size, overlap with ED≤ 2 frontier

### 2.6 Structural Analysis

#### 2.6.1 Conformer Ensemble Generation

For the top 5 evolved candidates and baselines, three-dimensional conformer ensembles were generated:

1. Use RDKit ETKDG (distance geometry with triangle bounds smoothing) to generate 8 diverse conformations for each unmodified peptide; for the lipidated Pal-KTTKS, where the long palmitoyl tail dominates the conformational extent, 3 conformations were used because additional samples added redundant rather than informative geometries.
2. Embed each conformation using distance-geometry distance bounds.
3. Optimize each structure with MMFF94s force field; fall back to UFF if MMFF fails.
4. Run 100 optimization cycles per conformer.
5. Compute radius of gyration (RoG) for each optimized structure.

##### Radius of Gyration

Defined as:

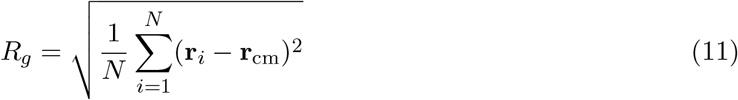

where **r**_*i*_ is the position of atom *i* and **r**_cm_ is the center of mass. RoG is a proxy for molecular size and flexibility; smaller RoG indicates a more compact, diffusion-favorable conformation.

##### Metrics reported

Mean, standard deviation, minimum, and maximum RoG across each conformer ensemble (8 conformers per unmodified peptide; 3 for Pal-KTTKS, consistent with the ensemble sizes described above).

### 2.7 Software and Reproducibility

All computations were performed using:

- Python 3.10.12
- RDKit 2023.09.1 (exact molecular descriptors)
- NumPy 1.24.3, Pandas 2.0.3, SciPy 1.11.1 (data handling)
- Custom implementations of GA, NSGA-II, Pareto utilities (no external MOEA libraries)

Random seeds were fixed (seed=42) to ensure deterministic reproducibility. All unit tests (48 tests covering descriptor calculations, constraint validation, optimization algorithms, and analysis) passed with 100% success rate. Complete code and results are available on GitHub: https://github.com/nkomianos/matrixyl-pareto-design.

## 3 Results

### 3.1 Tournament Search (Single-Objective)

Tournament selection with scalar fitness function *f* = penetration × function was run for 100 generations on a population of 100 individuals.

#### Best candidate

`PTTPS` (Pro-Thr-Thr-Pro-Ser)

- Optimization score: 0.4578
- Penetration score: 0.6866
- Functional preservation score: 0.6667
- Edit distance from KTTKS: 2 (mutations: K1P, K4P)
- Generation found: 4 (maintained as best through generation 100)

#### Descriptor profile for PTTPS

- Molecular weight: 501.54 Da (−11.0% vs KTTKS 563.65 Da)
- TPSA: 217.63 Å^2^ (−25.6% vs KTTKS 292.45 Å^2^)
- LogP: −3.98 (vs KTTKS −4.65; more balanced)
- H-bond donors: 8 (−27.3% vs KTTKS 11)
- H-bond acceptors: 9 (−18.2% vs KTTKS 11)
- Rotatable bonds: 11 (−45.0% vs KTTKS 20)
- Formal charge: 0 (vs KTTKS +2)

#### Convergence behaviour

The best candidate was found by generation 4 and remained optimal through all subsequent generations. Population mean fitness plateaued by generation 30. Early convergence of this kind is unsurprising on a 3,706-candidate search space: the optimizer quickly exhausts promising neighbours of the best individual and population diversity collapses.

### 3.2 Pareto Frontier (Multi-Objective)

NSGA-II identified a 9-member non-dominated frontier (Figure 1).

**Figure 1:**
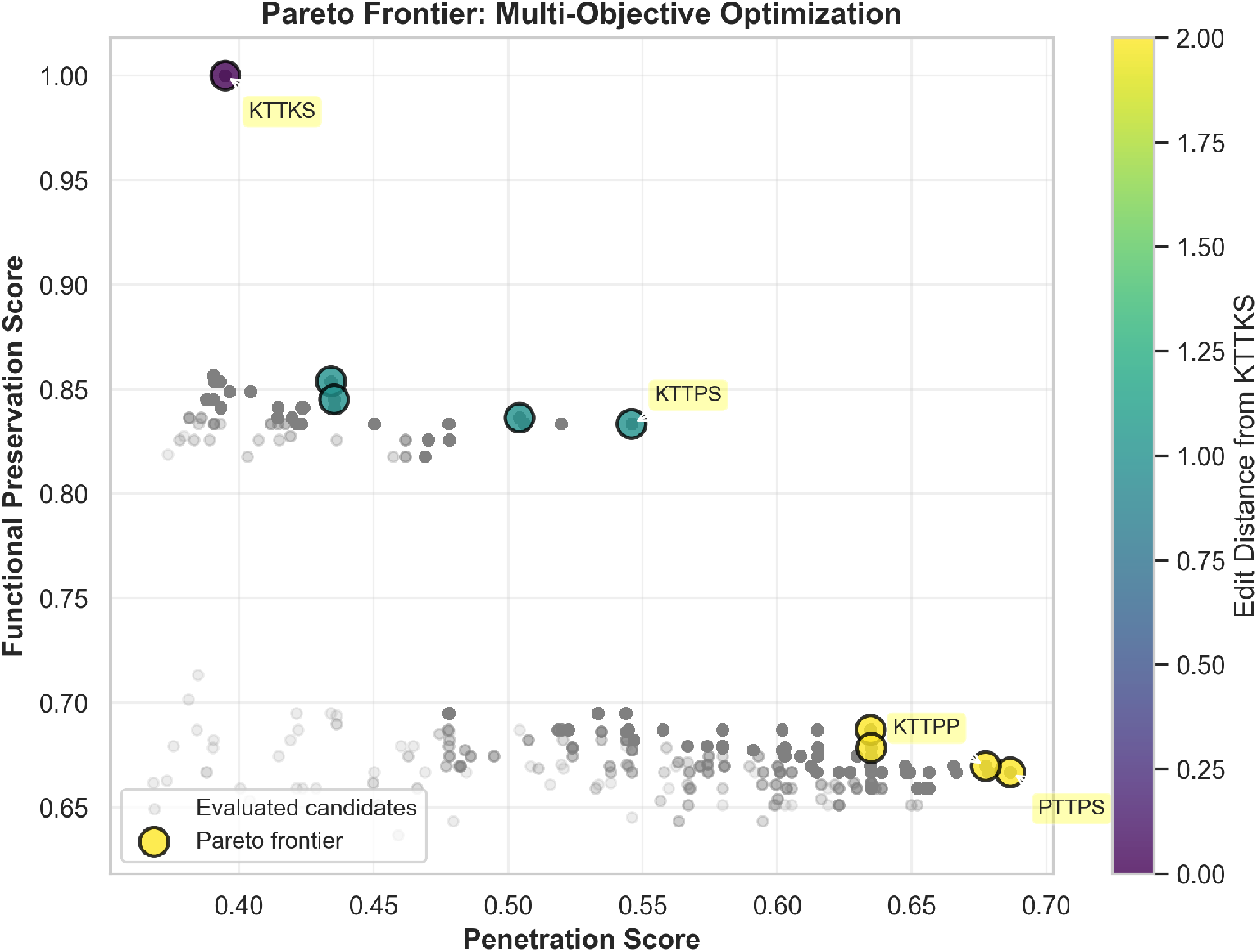
Pareto frontier from NSGA-II multi-objective optimization. Each point is a candidate peptide; the colored frontier members are non-dominated, meaning no other candidate is simultaneously better on all three objectives (penetration, functional preservation, synthesis feasibility). Both axes range over [0, 1]. All evaluated candidates are shown in gray; frontier members are colored by edit distance from the KTTKS baseline (viridis palette: dark = ED 0, bright = ED 2). Key candidates (PTTPS, KTTKS, KTTPS, KTTPP) are annotated.

#### Trade-off structure

1. **High-penetration zone** (penetration > 0.63): Candidates PTTPS, KTTPP, KTTPG, KTTPA cluster at edit distance 2 with penetration scores exceeding 0.63 but functional scores only 0.67–0.69.
2. **Balanced zone** (penetration 0.50–0.55): Candidates KTTKP, KTTPS occupy a middle ground with functional scores > 0.83.
3. **Function anchor** (baseline KTTKS): Perfect functional score (1.0) by definition but lowest penetration (0.395).

PTTPS and KTTPS occupy the two ends of the practical trade-off; KTTKS sits at the function-anchor end by definition.

Convergence behavior of both optimizers is summarized in Figure 2: the tournament GA reaches its best candidate by generation 4 and plateaus thereafter, while NSGA-II grows the non-dominated frontier monotonically to its 9-member final size.

**Figure 2:**
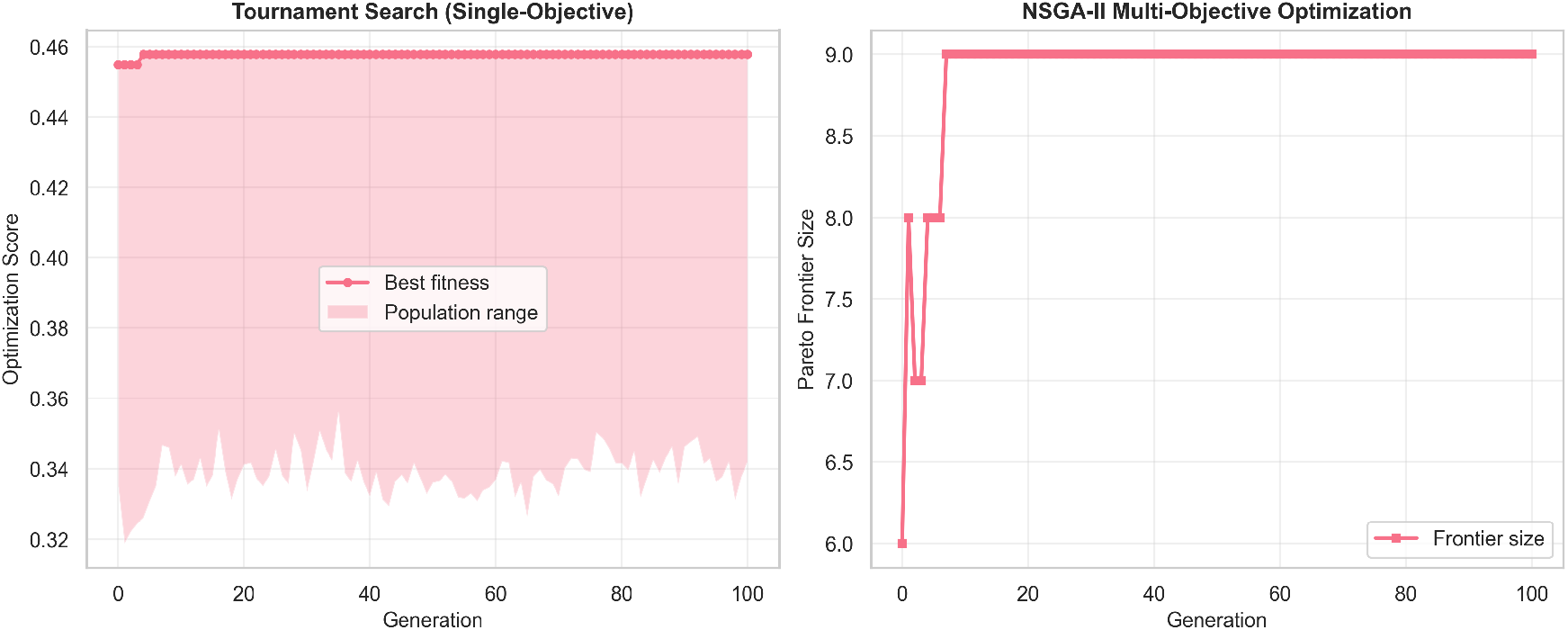
Convergence curves for optimization algorithms. Left: Tournament GA single-objective optimization showing best fitness and population range over 100 generations. Right: NSGA-II multi-objective optimization showing Pareto frontier size evolution. Both algorithms converge rapidly (best candidate found by generation 4 in tournament GA).

### 3.3 Mutation Enrichment Analysis

Across the 9-member frontier, only 5 unique mutations appear:

#### Interpretation

- **K4P dominance**: The K4P mutation is present in 5 of 9 frontier candidates (55.6%), far above the rate expected if mutations were drawn uniformly at random. Replacing lysine (polar, positively charged) with proline (hydrophobic, conformationally rigid) reduces TPSA, removes the +1 charge, and increases hydrophobic character. Each of these changes individually moves the candidate toward the descriptor ranges associated with passive transdermal diffusion.
- **Position 5 variability**: Serine at position 5 is replaced with three different residues (Ala, Gly, Pro) with nearly equal frequency (22.2% each). This suggests position 5 is less critical for permeability optimization and that multiple strategies (reducing H-bonds, adding flexibility, adding rigidity) are viable.
- **K1P rarity**: The N-terminal lysine K1P mutation is present only in PTTPS, the extreme high-penetration candidate. This suggests K1P provides additional benefit (charge reduction) but at higher cost to function recognition.

These position-by-mutation patterns are visualized in Figure 3.

**Figure 3:**
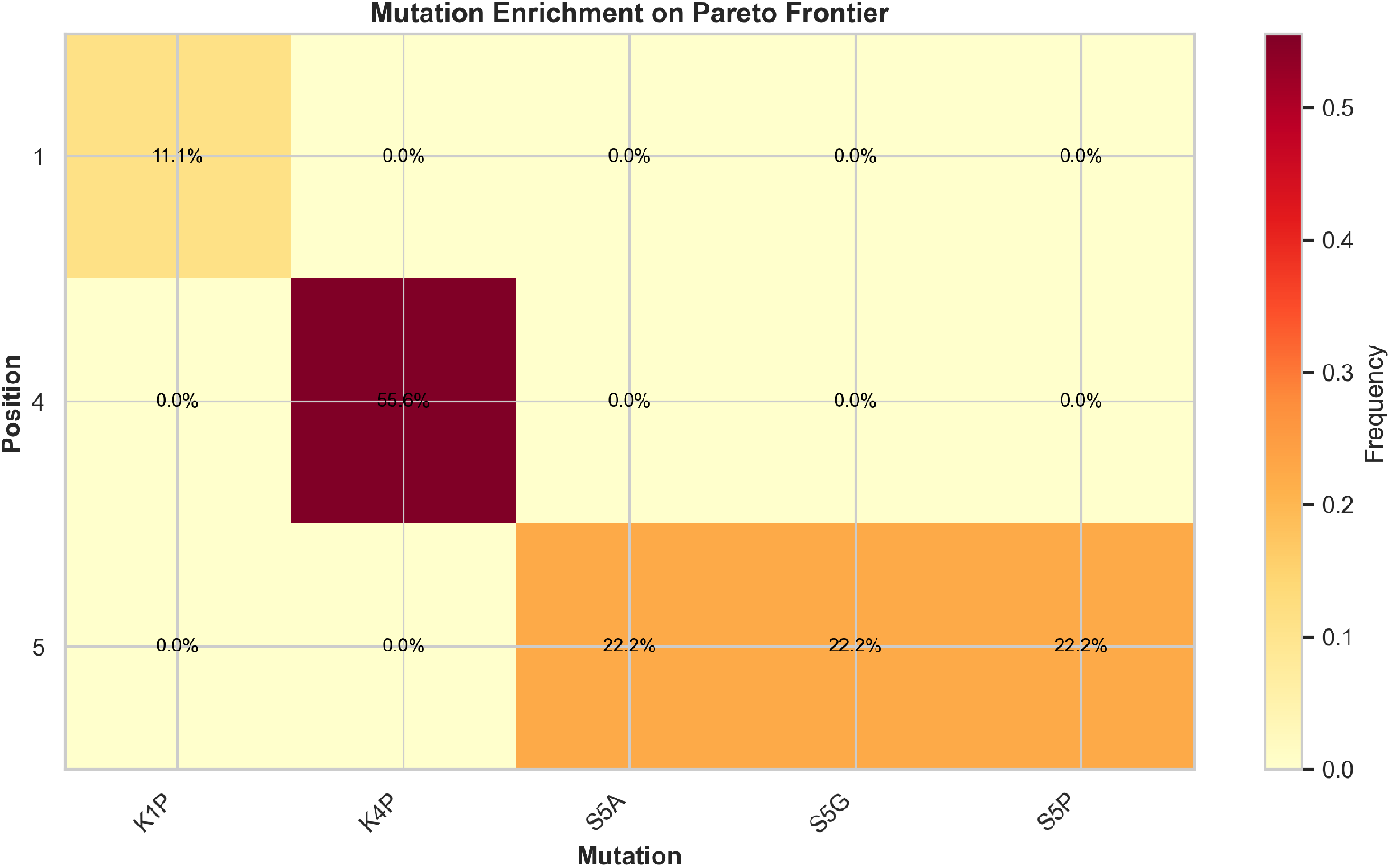
Mutation enrichment heatmap. Position (rows 1–5) versus amino-acid substitution (columns), showing the fraction of the 9-member Pareto frontier carrying each mutation (frequency = count / 9). K4P (Lys →Pro at position 4) is the dominant mutation, present in 55.6% of frontier candidates. Position 5 substitutions are more variable, with Ser →Ala, Ser →Gly, and Ser →Pro each at 22.2%.

### 3.4 Structural Validation (Conformer Ensembles)

RoG statistics for top candidates and baselines (8 conformers per sequence, MMFF94s/UFF optimization):

#### Key findings

1. **PTTPS is the most compact frontier member**: mean RoG of 5.05 Å is 6.9% smaller than the KTTKS baseline (5.42 Å), consistent with the hypothesis that K4P substitution (replacing the long, polar lysine side chain with the cyclic proline backbone) reduces conformational extent.
2. **Pal-KTTKS remains spatially extended**: although palmitoylation improves LogP by +0.99 against KTTKS, the palmitoyl tail extends the conformational ensemble to 8.38 Å mean RoG (54.5% larger than the unmodified peptide). The cost in molecular size therefore offsets the LogP benefit, which is why palmitoylation produces only a modest net gain in the penetration descriptor.
3. **Compactness is positively but imperfectly correlated with penetration**: the most compact candidate (PTTPS) also has the highest penetration score, and the high-penetration candidates cluster at the low-RoG end of the table. The ordering is not strictly monotonic, however: among the three high-penetration candidates the RoG ranking (PTTPS < KTTPS < KTTPP) differs from the penetration ranking (PTTPS > KTTPP > KTTPS), so RoG is one contributor to permeability rather than the sole driver.

### 3.5 Edit Distance 3 (Extended Search Space)

To test for diminishing returns, NSGA-II was re-run with max edit distance 3 (ED≤ 3, ≈ 100k candidates):

#### Interpretation

Extending to ED≤ 3 yields a slight penetration gain (0.785 vs 0.687, +14.2%), but:

1. The frontier expands from 9 to 20 candidates, reducing clarity of primary trade-off.
2. The new best candidate (KFFKP) involves two phenylalanine substitutions, moving significantly away from the recognizable Matrixyl motif.
3. All original ED≤ 2 frontier members remain on the ED≤ 3 frontier, indicating ED≤ 2 captures the essential trade-off structure.

For this manuscript, ED≤ 2 is preferred: it provides clearer narrative, maintains closer motif proximity, and reduces candidate burden for experimental validation.

Figure 4 situates the Pareto frontier within the full 3,706-candidate evaluated search space, and Figure 5 shows the edit-distance composition of the frontier relative to that space.

**Figure 4:**
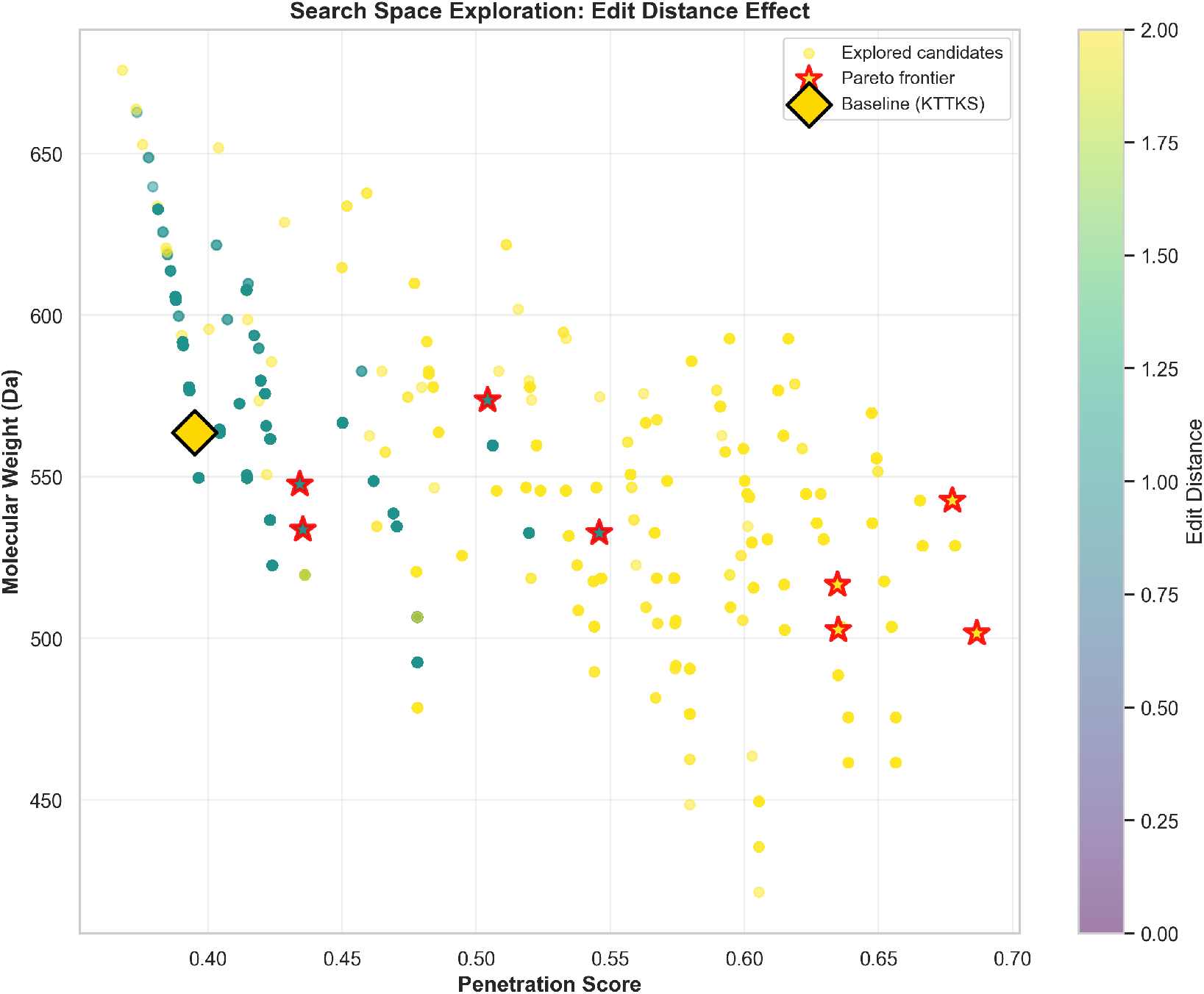
Search space visualization. Each point is one of the 3,706 candidates in the ED ≤2 neighborhood of KTTKS, plotted as penetration score (*x*-axis) against molecular weight (*y*-axis) and colored by edit distance from the baseline (viridis palette: dark = ED 0, bright = ED 2). Pareto frontier members are starred; baseline KTTKS is shown as a gold diamond. The frontier dominates the high-penetration, low-*M*_*w*_ region of the search space.

**Figure 5:**
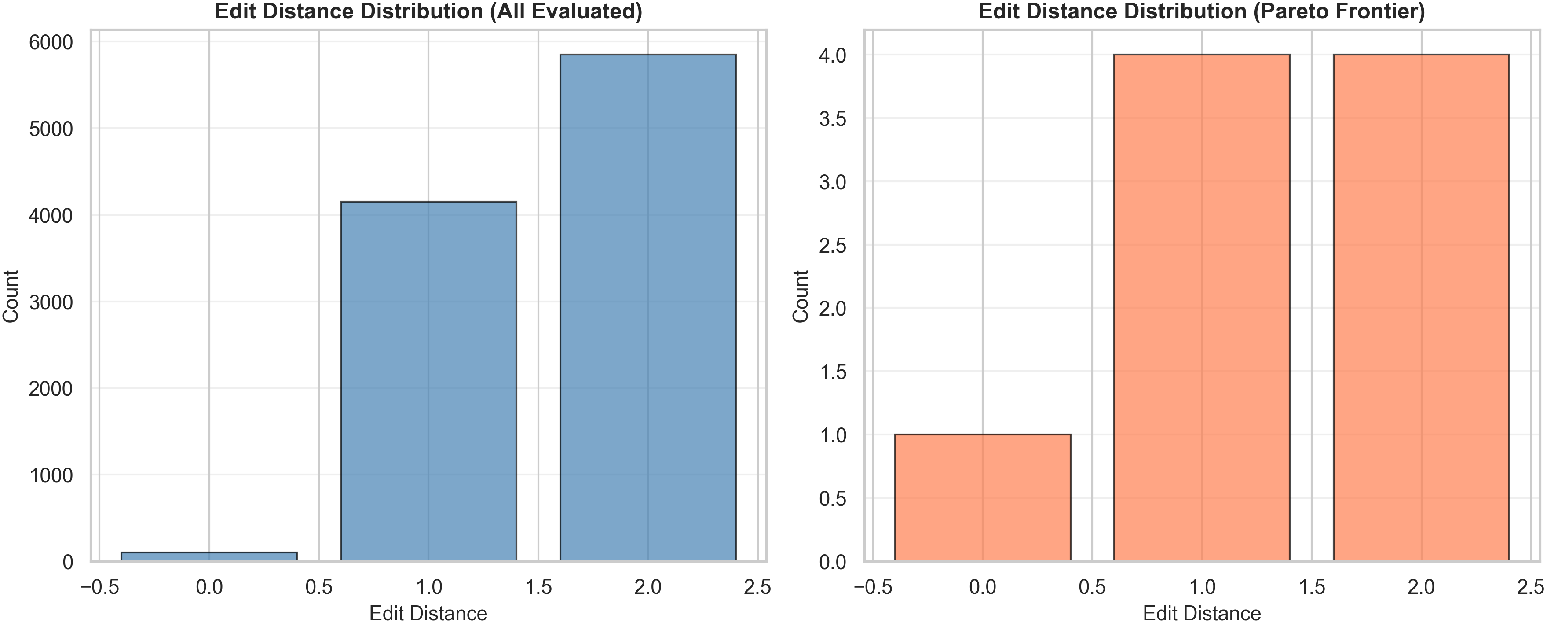
Edit distance distribution. Left: full enumerated search space showing the edit-distance histogram (ED = 0: 1 baseline, ED = 1: 95 candidates, ED = 2: 3,610 candidates; total 3,706). Right: Pareto frontier composition by edit distance (majority at ED = 1, 2, with the baseline at ED = 0). The frontier captures candidates at every edit-distance class within the ED ≤ 2 neighbourhood.

### 3.6 Sensitivity Analysis (Parameter Robustness)

±30% variations in TPSA, *M*_*w*_, and LogP penalties were applied to the Phase 2 frontier. Results:

#### Conclusion

The frontier ranking is completely stable under five of the six perturbations: ±30% changes to the TPSA or *M*_*w*_ penalties, and a 30% increase in the LogP penalty, all leave every candidate’s position unchanged. The single exception is a 30% reduction of the LogP penalty, which reorders 6 of 9 frontier members and demotes the top candidate. This is consistent with LogP being the only descriptor that is far from its target range for several frontier members; reducing its penalty allows otherwise-suppressed hydrophilic candidates to re-enter the top. The robust K4P signature and the high-penetration core of the frontier (PTTPS, KTTPP) are not artifacts of the specific weighting we chose; they survive all ±30% perturbations of the TPSA and *M*_*w*_ penalties.

### 3.7 Exhaustive Enumeration Validation

The full 3,706-candidate space was exhaustively evaluated and ranked by Pareto dominance:

#### Interpretation

NSGA-II with the parameters described in Section 2.4.2 recovered the full ground-truth Pareto set on this 3,706-candidate space. For spaces of this size and structure, the evolutionary search is sufficient; for larger spaces where exhaustive enumeration is infeasible, the same algorithm would have no such guarantee.

## 4 Discussion

### 4.1 Interpretation of Results

#### 4.1.1 K4P at position 4

Five of nine frontier members carry the K4P substitution, far more than would be expected from uniform-random sampling. The position-4 lysine in KTTKS is conserved across matrikines and has been argued to participate in electrostatic interactions with collagen [13], so its replacement is not innocuous from the standpoint of biological activity. From the standpoint of the descriptor scores it is straightforward: lysine to proline removes one unit of positive charge, increases hydrophobicity (Kyte-Doolittle index +1.8 for proline versus −3.9 for lysine), and replaces a hydrogen-bond donor with a residue that has none. Each of these changes individually moves the candidate toward the Potts and Guy ranges for passive transdermal diffusion.

There is a corresponding cost on the functional side. The functional-preservation score for the leading K4P-bearing candidate, PTTPS, is 0.67 against a baseline of 1.00 for KTTKS. The fact that K4P-bearing candidates remain on the frontier even after this penalty, rather than being dominated by candidates that retain the lysine, is the meaning of “Pareto-optimal” in this context: any further improvement in the penetration descriptor requires losing more functional preservation than K4P already costs.

#### 4.1.2 Position 5 Flexibility

Position 5 (serine in KTTKS) shows variable substitution patterns: Ala, Gly, and Pro each appear in 2 frontier candidates (22.2% frequency each). This heterogeneity suggests that position 5 is less constrained than position 4. Reducing hydrogen-bond donors via alanine, adding backbone flexibility via glycine, and restricting backbone via proline all yield comparable permeability gains. This is consistent with position 5 being distal from the known functional epitope of KTTKS.

#### 4.1.3 Three regions of the frontier

For experimental triage, the natural primary candidate is KTTPS, the single-mutation analog: it has the highest prior probability of retaining biological activity (motif-preservation 0.833) while still gaining 38% in the penetration descriptor against KTTKS. PTTPS is the appropriate aggressive arm for testing whether descriptor gains at the limit translate into measurable permeation, accepting a larger functional cost. KTTPP serves as a synthesis backup with a profile close to PTTPS but a slightly higher functional score. KTTKS is the unmodified control.

### 4.2 Comparison to Prior Work

#### 4.2.1 Protein Language Models

PLM-based peptide and protein design [8] optimizes for sequence naturalness, that is, sequences that look plausible to a model trained on the protein universe. The objective in this paper is different: we optimize for non-evolutionary, chemistry-derived descriptors of passive diffusion (LogP near 2, TPSA below 140 Å^2^, *M*_*w*_ below 500 Da) that have no direct counterpart in the training distribution of a PLM. The two objectives can be combined. PLM embeddings could enter the multi-objective fitness as an additional naturalness term alongside the RDKit descriptors used here, with the Pareto frontier then quantifying the trade-off between naturalness and predicted delivery.

#### 4.2.2 Transdermal Delivery Literature

Our descriptor thresholds (*M*_*w*_ < 500, TPSA < 140) derive from Potts & Guy (1992) [4] and Prausnitz, Mitragotri & Langer (2004) [12], which synthesized transdermal permeability data from hundreds of small molecules. Applying these thresholds to peptides is a deliberate extrapolation, justified by the observation that peptides generally follow similar rules but with higher variance [3]. By using soft penalties rather than hard cutoffs, we acknowledge this uncertainty while still guiding optimization toward delivery-favorable ranges.

#### 4.2.3 Computational peptide design

Computational peptide design has typically combined evolutionary or generative search with learned scoring functions trained on a limited amount of experimental data. The work presented here makes a different set of choices. The scoring is done with exact RDKit molecular descriptors rather than learned proxies. The optimization is multi-objective (a Pareto frontier) rather than a single scalarized fitness, so the trade-off between objectives is visible directly. The frontier is checked against the ground-truth Pareto set obtained by exhaustive enumeration of the search space, rather than against an approximate benchmark. Each candidate on the frontier can be explained in terms of changes to specific descriptors, so the output of the search is a small set of testable hypotheses about how individual substitutions move the candidate through descriptor space.

### 4.3 Limitations and Future Work

#### 4.3.1 No Receptor-Level Model

Our functional-preservation score uses sequence motif similarity, not a validated collagen-binding or fibroblast-activation assay. While the BLOSUM62 scoring incorporates biochemical knowledge of amino-acid conservation, we make no claim that K4P substitution retains actual collagen-stimulating activity. Experimental validation (fibroblast collagen secretion assays) is essential before declaring candidates “functional.”

#### 4.3.2 Conformers as Exploratory Metrics

RDKit conformer ensembles provide useful proxies for molecular size and flexibility but are not equivalent to full molecular dynamics simulations. The MMFF94s force field does not explicitly model aqueous environments, membrane interactions, or kinetic barriers. We use RoG correlations as suggestive evidence (supporting the diffusion hypothesis) rather than proof.

#### 4.3.3 No Skin-Specific Binding or Metabolism

The penetration score penalizes TPSA and *M*_*w*_ globally; it does not model stratum corneum-specific protein binding, enzymatic degradation, or pH-dependent charge states in different skin compartments. These factors could significantly modulate actual permeability and are left to experimental determination.

#### 4.3.4 Edit Distance 2 is Conservative

Restricting to ED ≤ 2 maintains motif proximity but may exclude superior candidates at ED = 3 or higher. Our ED = 3 analysis shows only modest improvement (0.687 → 0.785, +14.2%), but full space exploration is infeasible for larger problems. Future work should apply the same framework to less-constrained spaces (variable-length sequences, non-canonical residues) where exhaustive validation is impossible and external validation (e.g., PLM uncertainty quantification) becomes more important.

#### 4.3.5 Future Directions

1. **Hybrid PLM + chemical optimization**: combine PLM-derived naturalness scores (for example from ESM-2 or a ProGen-style decoder model) with the RDKit delivery descriptors used here in a single multi-objective fitness.
2. **Active learning from experiments**: Run small batches of in vitro permeation and fibroblast assays, use results to refine functional-preservation scoring, re-optimize.
3. **Formulation co-optimization**: Extend the framework to jointly optimize peptide sequence and formulation parameters (penetration enhancer concentration, vehicle composition).
4. **Larger peptides**: Apply the same methodology to longer cosmetic or therapeutic peptides (10–20 residues), where exhaustive enumeration is infeasible but MOEA + sensitivity analysis remain tractable.

## 5 Conclusions

The Pareto frontier in the trade-off between predicted skin permeability and motif preservation for the Matrixyl core peptide KTTKS has nine members, and is identical to the ground-truth set recovered by exhaustive enumeration of the 3,706-candidate edit-distance-2 neighbourhood. Lysine-to-proline at position 4 (K4P) is the single substitution that dominates this frontier: it appears in five of nine frontier members, recurs across all ±30% perturbations to the TPSA and *M*_*w*_ penalty weights, and can be read directly in terms of charge, polarity, and hydrogen-bond count. We propose KTTPS as the primary candidate for experimental validation (single mutation, motif-preservation 0.833, 38% gain in the penetration descriptor), PTTPS as the aggressive arm (two mutations, 74% gain at a functional cost), and KTTPP as a synthesis backup. All code, results, and figures are openly available; the 48-test unit-test suite covers the descriptor calculations, the search-space rules, and the optimizer implementations.

## Author Contributions

N.K. and P.P. contributed equally to this work. Per the CRediT taxonomy, both authors share the following roles: **Conceptualization, Methodology, Software, Validation, Formal analysis, Investigation, Data curation, Visualization, Writing – original draft, Writing – review & editing**, and **Project administration**. Both authors read and approved the final manuscript.

## Ethics

This study did not involve human or animal subjects. All work is computational and relies exclusively on publicly available reference data (PubChem CID 9897237 for palmitoyl pentapeptide-4; canonical amino-acid sequence libraries).

## Use of Generative AI

In line with bioRxiv’s policy on disclosure of artificial intelligence use, the authors note the following. The scientific design of the work, all algorithmic choices, all hyperparameters, all interpretations of the results, and all written claims in this manuscript are the responsibility of the human authors. Large language models (Anthropic Claude) were used in two capacities during the project. First, as an editorial assistant during drafting of the manuscript and the supporting documentation in the public repository. Second, as a code-completion tool during implementation of the pipeline; every routine was read, tested, and signed off by a human author, and the 48-test unit-test suite was written and reviewed by the human authors. No part of the manuscript was written or accepted without human review. No scientific claim, numerical result, or citation in this manuscript was generated solely by an AI system without human verification. No generative AI is listed as an author of this work, and no AI system holds responsibility for the content.

## Data and Code Availability

All source code, unit tests, per-phase result CSVs, and publication figures are openly available at https://github.com/nkomianos/matrixyl-pareto-design. A persistent snapshot of the release accompanying this preprint is archived on Zenodo with DOI 10.5281/zenodo.20128083 (git tag v1.0.1-preprint, commit fae1b56). The repository is licensed under MIT (code, data, results); this preprint is distributed under the Creative Commons Attribution 4.0 International License (CC BY 4.0). No proprietary or restricted data are used in this study.

## A Supplementary Methods

### A.1 BLOSUM62 Scoring Details

The BLOSUM62 matrix (Blocks Substitution Matrix, derived from aligned protein blocks) provides log-odds scores for amino-acid substitutions observed in nature at ≥ 62% sequence identity. A positive score indicates a substitution is more frequent than expected by chance (conservative), while a negative score indicates it is rare (non-conservative).

For position-by-position functional-preservation scoring, we normalized BLOSUM62 scores by dividing by 4.0 (the maximum BLOSUM62 value, for identical residues) and clamping to [0, 1]:

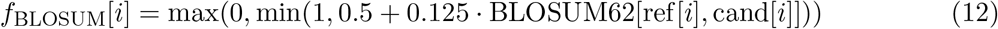

This maps BLOSUM62 scores in the range [−4, +4] to [0.5, 1], ensuring even mismatch-penalizing (negative BLOSUM62) substitutions receive some credit if they are less bad than random.

### A.2 Search Space Enumeration

The 3,706-candidate fixed-length ED≤ 2 neighborhood was generated via dynamic programming:

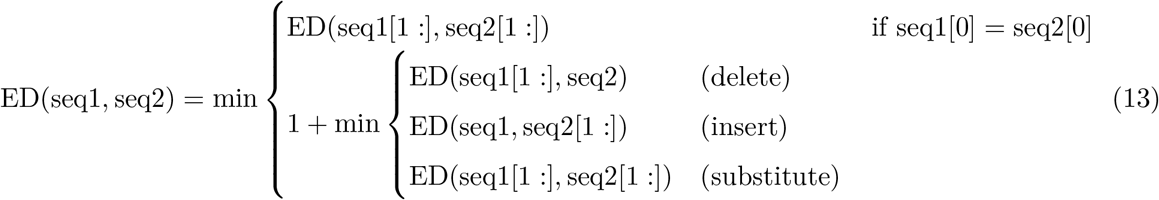

For fixed-length sequences of length 5, insertions and deletions are not applicable, so ED reduces to the number of substitutions required (Hamming distance for equal-length strings). The enumeration algorithm:

1. Iterate over all 20 canonical amino acids at position 1: 20 candidates.
2. For each, iterate over all 20 at position 2: 20 × 20 candidates.
3. Continue for positions 3, 4, 5.
4. For each 5-residue sequence, compute Hamming distance to KTTKS.
5. Retain only sequences with Hamming distance ≤ 2.

This yields:

- ED = 0: 1 candidate (KTTKS itself)
- 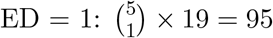 candidates (5 positions, 19 alternative residues per position)
- 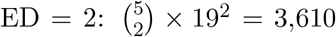 candidates (two positions changed, each replaced by one of 19 alternatives)
- Total: 1 + 95 + 3,610 = 3,706 candidates

This count is also recovered by enumerating all 20^5^ length-5 sequences and post-filtering for Hamming distance ≤ 2, which we use as an independent check.

### A.3 Code Availability

All source code, unit tests, results CSVs, and figures are available at:

https://github.com/nkomianos/matrixyl-pareto-design

Unit test suite (48 tests):

`python -m pytest tests/ -v`

Phase 1 execution:

~~~
python experiments/01_tournament_search.py \
 --sequence data/sequences/matrixyl_core.fasta \
 --population 100 --generations 100 --seed 42 \
 --output results/phase1_tournament
~~~

Phase 2 execution:

~~~
python experiments/02_nsga2_pareto.py \
 --sequence data/sequences/matrixyl_core.fasta \
 --population 100 --generations 100 --seed 42 \
 --output results/phase2_pareto
~~~

All commands logged with timestamps and git commit information in run_metadata.json within each output directory.

## References

[1] Henninot, A., Collins, J. C., Nuss, J. M. The current state of peptide drug discovery: back to the future? J. Med. Chem. 2018, 61, 1382–1414. doi:10.1021/acs.jmedchem.7b00318.

[2] Robinson, L. R., Fitzgerald, N. C., Doughty, D. G., Dawes, N. C., Berge, C. A., Bissett, D. L. Topical palmitoyl pentapeptide provides improvement in photoaged human facial skin. Int. J. Cosmet. Sci. 2005, 27, 155–160. doi:10.1111/j.1467-2494.2005.00261.x.

[3] Mitragotri, S., Burke, P. A., Langer, R. Overcoming the challenges in administering biopharmaceuticals: formulation and delivery strategies. Nat. Rev. Drug Discov. 2014, 13, 655–672. doi:10.1038/nrd4363.

[4] Potts, R. O., Guy, R. H. Predicting skin permeability. Pharm. Res. 1992, 9, 663–669. doi:10.1023/A:1015810312465.

[5] Lintner, K., Peschard, O. Biologically active peptides: from a laboratory bench curiosity to a functional skin care product. Int. J. Cosmet. Sci. 2000, 22, 207–218. doi:10.1046/j.1467-2494.2000.00010.x.

[6] Lin, Z., Akin, H., Rao, R., Hie, B., Zhu, Z., Lu, W., et al. Evolutionary-scale prediction of atomic-level protein structure with a language model. Science 2023, 379, 1123–1130. doi:10.1126/science.ade2574.

[7] Jumper, J., Evans, R., Pritzel, A., Green, T., Figurnov, M., Ronneberger, O., et al. Highly accurate protein structure prediction with AlphaFold. Nature 2021, 596, 583–589. doi:10.1038/s41586-021-03819-2.

[8] Madani, A., Krause, B., Greene, E. R., Subramanian, S., Mohr, B. P., Holton, J. M., et al. Large language models generate functional protein sequences across diverse families. Nat. Biotechnol. 2023, 41, 1099–1106. doi:10.1038/s41587-022-01618-2.

[9] Deb, K., Pratap, A., Agarwal, S., Meyarivan, T. A fast and elitist multiobjective genetic algorithm: NSGA-II. IEEE Trans. Evol. Comput. 2002, 6, 182–197. doi:10.1109/4235.996017.

[10] Francoeur, P. G., Masuda, T., Sunseri, J., Jia, A., Iovanisci, R. B., Snyder, I., Koes, D. R. Three-dimensional convolutional neural networks and a cross-docked data set for structure-based drug design. J. Chem. Inf. Model. 2020, 60, 4200–4215. doi:10.1021/acs.jcim.0c00411.

[11] Watson, J. L., Juergens, D., Bennett, N. R., Trippe, B. L., Yim, J., Eisenach, H. E., et al. De novo design of protein structure and function with RFdiffusion. Nature 2023, 620, 1089–1100. doi:10.1038/s41586-023-06415-8.

[12] Prausnitz, M. R., Mitragotri, S., Langer, R. Current status and future potential of transdermal drug delivery. Nat. Rev. Drug Discov. 2004, 3, 115–124. doi:10.1038/nrd1304.

[13] Katayama, K., Armendariz-Borunda, J., Raghow, R., Kang, A. H., Seyer, J. M. A pentapeptide from type I procollagen promotes extracellular matrix production. J. Biol. Chem. 1993, 268, 9941–9944. doi:10.1016/S0021-9258(18)82153-6.

